# Caspase-7 cleavage repurposes AAK1 to suppress mRNA decay and promote tolerance to anti-tubulin agents

**DOI:** 10.64898/2026.11.20.767991

**Authors:** Yunhan Jiang, Kunyang Li, Pawel Nowialis, Exing Wang, Zhao Zhang, Xiaobo Mao, Nu Zhang, Gang Huang, Lu-Zhe Sun, Ratna Vadlamudi, Zhijian Qian, Guangrong Zheng, Shuang Huang, Lina Song, Sumin Kang, Lingtao Jin

## Abstract

Prolonged mitotic arrest imposed by anti-tubulin chemotherapy suppresses transcription, making mRNA preservation critical for tumor-cell survival. Here, we identify an unexpected kinase-repurposing mechanism that sustains this adaptation. A kinome-wide screen revealed clathrin-adaptor kinase AAK1 as a determinant of taxane tolerance. During mitotic arrest, CDK1 phosphorylates AAK1 at T389 to enhance catalytic activity, while caspase-7 cleaves AAK1 at D767, removing a clathrin-binding domain. These modifications produce a cleaved fragment with reduced clathrin association and increased association with CCR4–NOT, suppressing mRNA decay and maintaining glycolytic gene expression. Mutant-rescues establish that both phosphorylation and cleavage contribute to cancer-cell survival during paclitaxel-induced arrest. A clinical-stage AAK1 inhibitor developed for neuropathic pain sensitized cancer cells to paclitaxel. Together, these findings reveal how coordinated phosphorylation and proteolysis repurpose a trafficking kinase to maintain mRNA stability, uncovering an unexpected tolerance-promoting function of caspase cleavage and a therapeutically actionable vulnerability of chemotherapy-arrested cancer cells.

## Background

Anti-tubulin agents, including the taxanes paclitaxel and docetaxel and the vinca alkaloids, remain among the most widely used chemotherapeutics for ovarian, breast, non-small cell lung, and head and neck cancer, as well as for lymphoma and leukemia. In ovarian cancer, paclitaxel/platinum combination therapy has been the cornerstone of first-line treatment for more than three decades and produces high initial response rates. Nevertheless, most patients with advanced disease ultimately relapse with chemoresistant disease, which remains the principal cause of mortality^1,2^. Resistance has been attributed to altered microtubule dynamics, spindle checkpoint adaptation, and drug efflux, but the signaling pathways that allow cancer cells to survive prolonged mitotic arrest are largely undefined. Arrested-cell fate reflects competition between accumulating pro-apoptotic signals and cyclin B degradation that permits mitotic slippage. Targeting signaling that shifts this balance could enhance anti-tubulin agent response.

During prolonged mitotic arrest, transcription is globally suppressed, cap-dependent translation is attenuated, and cells must sustain ATP production. Transcript abundance therefore depends increasingly on mRNA decay. The principal effector of that machinery is CCR4-NOT, a multisubunit complex that shortens poly(A) tails through its CNOT6/CNOT6L and CNOT7/CNOT8 nuclease modules and thereby commits transcripts to degradation^3^. Importantly, recent work showed that prolonged mitotic arrest globally stabilizes mRNAs, with poly(A)-tail profiles consistent with reduced deadenylation and stabilization dependent on PABPC1 and PABPC4^4^. Arrested cells require ATP for chromosome cohesion, spindle maintenance and membrane integrity, and glycolytic flux has been reported to support survival under these conditions^5,6^. How mitotic stress signaling is communicated to the decay machinery, and whether protein kinases participate directly in remodeling RNA regulatory complexes during chemotherapy, remain largely unexplored.

A second feature of prolonged mitotic arrest is engagement of the apoptotic protease machinery. Executioner caspase activity, however, is not an all-or-none commitment to death. Sublethal caspase activation has established non-apoptotic roles in neuronal remodeling, differentiation, and tissue morphogenesis, where limited proteolysis reshapes substrate function rather than destroying it. Caspase cleavage can activate a substrate, relocalize it, or remove a regulatory domain to expose an otherwise cryptic interaction surface, and proteome-wide surveys indicate that most caspase substrates are cleaved at a single site rather than degraded^7,8^. How caspase-mediated remodeling of existing proteins contributes to mRNA preservation and taxane tolerance during mitotic arrest remains poorly understood.

To identify kinase pathways required for chemotherapy resistance, we established a kinome-wide functional RNAi screening platform targeting 781 human kinases, an approach that has previously uncovered signaling mechanisms governing platinum resistance^9,10^. The paclitaxel screen identified AP2-associated kinase 1 (AAK1). AAK1 is one of four numb-associated kinases, alongside GAK, BIKE/BMP2K and STK16, and regulates clathrin-mediated endocytosis. It phosphorylates the AP2 mu2 subunit to promote cargo recruitment, and two tandem clathrin-binding domains in its C-terminal half anchor it to assembling coated pits^11,12^. Its roles in paclitaxel response and RNA metabolism remain unexplored. Separately, genetic and pharmacological inhibition of AAK1 relieves neuropathic pain, and selective AAK1 inhibitors have advanced into clinical trials for that indication^13–15^. This is a notable convergence, since peripheral neuropathy is the dose-limiting toxicity of the taxanes, and it provides a rationale for evaluating AAK1 inhibition in taxane-treated tumors.

Here we show that caspase-7-mediated cleavage repurposes AAK1 during chemotherapy-induced mitotic arrest. Cleavage at D767 removes the second clathrin-binding domain and generates a fragment with reduced clathrin association and enhanced association with CCR4–NOT, while CDK1 phosphorylation at T389 enhances catalytic activity. Together, these changes link AAK1 to suppression of mRNA decay, maintenance of glycolytic gene expression and paclitaxel tolerance, providing a preclinical rationale for targeting this dependency.

## Results

### Kinome-wide RNAi screening identifies AAK1 as a synthetic lethal partner for anti-tubulin agents

To systematically identify kinases dictating anti-tubulin sensitivity, we performed kinome-wide RNAi screening using a lentiviral shRNA library targeting 781 human kinase and kinase-related genes (4,518 clones). Paclitaxel-resistant KB-3-1^TaxolR^ cells were transduced with the lentivirus pool and treated with sublethal doses of paclitaxel. Among the top candidates, we identified three functional groups: established cell cycle and microtubule regulators (Aurora B, MARK1, CCND1) known to influence paclitaxel sensitivity^16,17^; kinases in TGF-β signaling (BMPR1A, BMPR2, ACVR1C) previously implicated in paclitaxel resistance^18^; and AAK1, which had no known connection to paclitaxel sensitivity (Figure 1A). Given the unclear function of AAK1 in this context, we focused subsequent investigations on this kinase.

**Figure 1.**
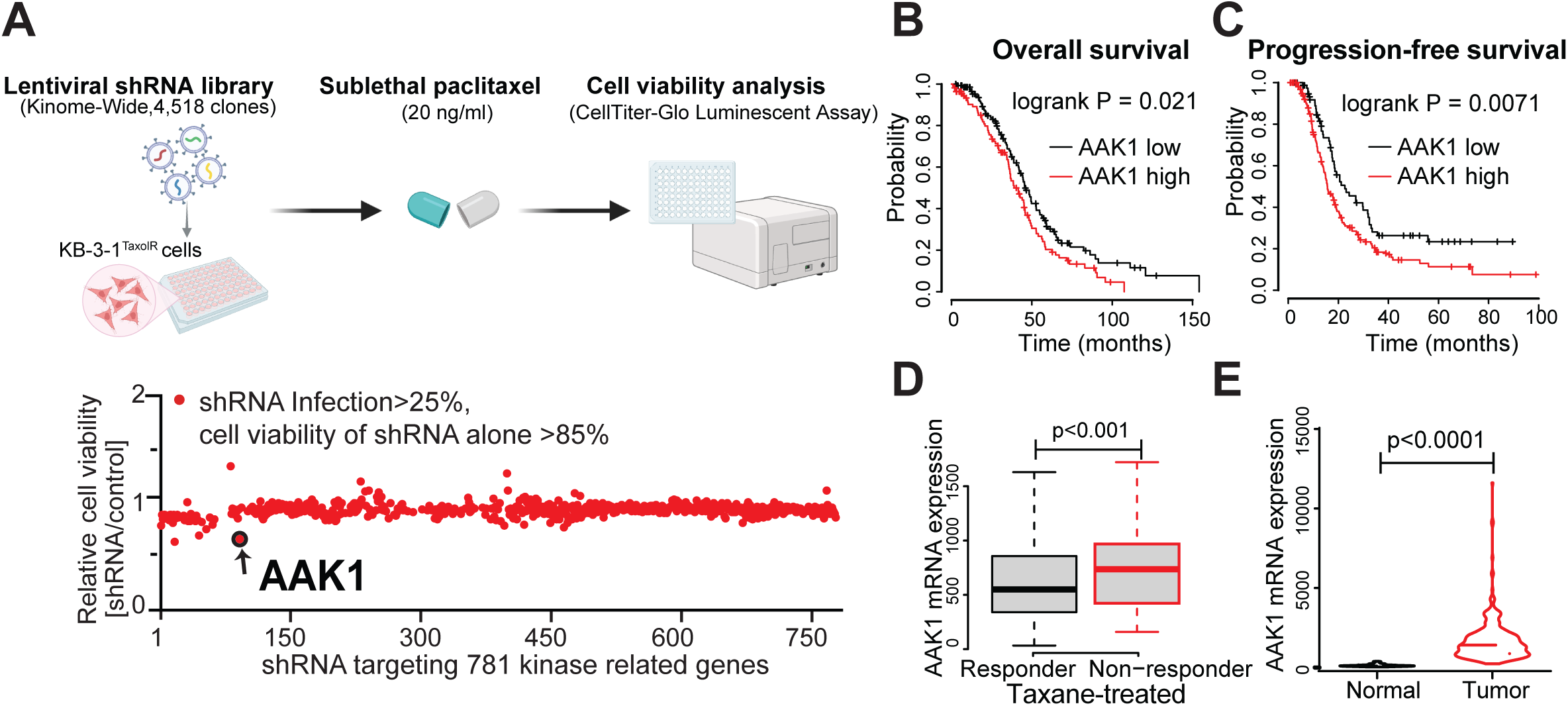
Kinome-wide RNAi screening identifies AAK1 as a synthetic lethal partner for paclitaxel in ovarian cancer. (A) Schematic of the kinome-wide shRNA screen in paclitaxel-resistant KB-3-1TaxolR cells (4,518 clones targeting 781 kinase genes; 20 ng/mL paclitaxel; CellTiter-Glo readout) and relative cell viability (shRNA/control) for each clone passing the infection (>25%) and single-agent viability (>85%) filters, with AAK1 indicated. (B–C) Kaplan-Meier analysis of overall survival (B) and progression-free survival (C) in taxane-treated ovarian cancer patients stratified by AAK1 expression (Kmplot). (D) AAK1 mRNA expression in taxane-treated responders versus non-responders (ROCplot, pooled ovarian cancer datasets). (E) AAK1 mRNA expression in ovarian tumors versus normal tissue (ROCplot).

To identify the cancer type in which AAK1 may play the most significant role in paclitaxel sensitivity, we analyzed TCGA datasets of paclitaxel-treated patient populations. High AAK1 expression was significantly correlated with both poor overall survival (Figure 1B) and progression-free survival (Figure 1C) in ovarian cancer patients. ROCplot analysis of pooled ovarian cancer patient datasets further revealed that paclitaxel non-responders exhibited higher AAK1 levels compared to responders^19,20^ (Figure 1D). AAK1 expression was also elevated in ovarian tumors compared to normal tissues (Figure 1E).

### AAK1 kinase activity promotes paclitaxel resistance beyond its canonical endocytosis role

To examine whether AAK1 regulates cellular response to paclitaxel, we silenced AAK1 in ovarian cancer cell lines and patient-derived organoids. AAK1 knockdown using shRNA significantly sensitized cells to paclitaxel (Figures 2A–2D) and docetaxel (Figure 2E) across all models tested. Importantly, AAK1 silencing sensitized cells to vinca alkaloids (vincristine) but not to platinum agents (Figures 2F–2G), indicating a specific role for AAK1 in regulating sensitivity to anti-tubulin agents. Mechanistically, AAK1 knockdown enhanced mitotic arrest and apoptosis in the presence of paclitaxel (Figures 2H–2I). *In vivo* xenograft assays further demonstrated that CRISPR-mediated AAK1 depletion sensitized ovarian cancer xenografts to paclitaxel (Figure 2J).

**Figure 2.**
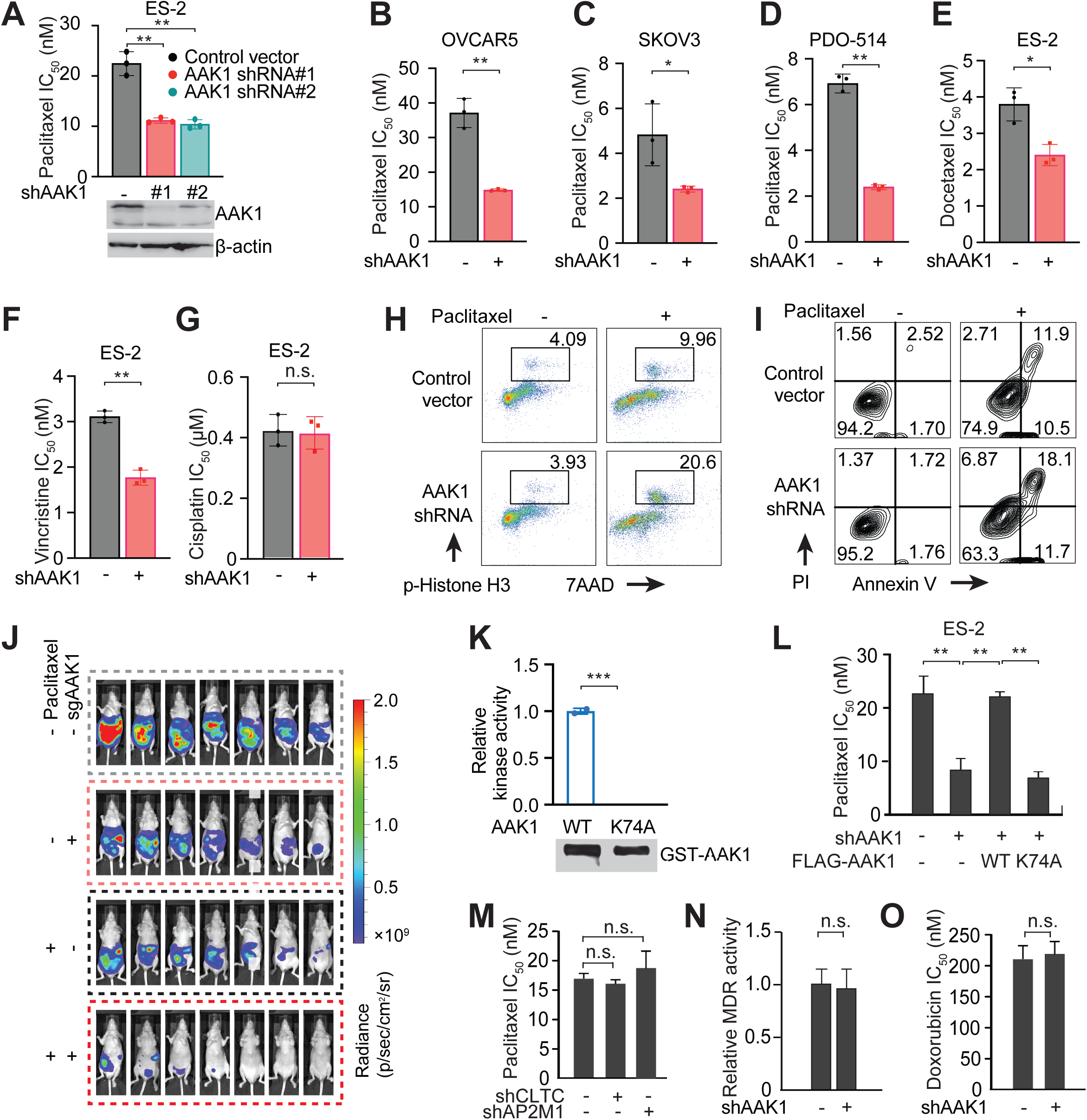
AAK1 kinase activity promotes paclitaxel resistance beyond its canonical endocytosis role. (A–D) Paclitaxel IC50 in ES-2 cells expressing two independent AAK1 shRNAs, with immunoblot confirmation of knockdown (A), and in OVCAR5 (B), SKOV3 (C) and the patient-derived organoid PDO-514 (D). (E–G) IC50 of ES-2 cells for docetaxel (E), vincristine (F) and cisplatin (G) with or without AAK1 knockdown. (H) Cell cycle analysis (p-Histone H3/7AAD) and (I) Annexin V/PI apoptosis assay in control and AAK1 knockdown ES-2 cells with or without paclitaxel. (J) Bioluminescence imaging of ES-2 xenografts treated with vehicle or paclitaxel, with or without sgAAK1 (n = 7 per group). (K) *In vitro* kinase activity of AAK1 wild-type versus kinase-dead K74A, with GST-AAK1 input. (L) Paclitaxel IC50 in AAK1 knockdown ES-2 cells reconstituted with shRNA-resistant FLAG-AAK1 wild-type or K74A. (M) Paclitaxel IC50 after knockdown of clathrin heavy chain (CLTC) or AP2M1. (N) MDR activity and (O) doxorubicin IC50 in control versus AAK1 knockdown cells. Data are mean ± SD, n = 3 biological replicates. *p < 0.05, **p < 0.01, n.s. not significant; two-tailed Student’s t-test. (A-I, K-O) Results of one representative experiment from 2-3 independent experiments are shown.

To determine whether kinase activity is required, we generated shRNA-resistant AAK1 wild-type and a kinase-dead mutant (K74A) and confirmed that K74A abolishes catalytic activity *in vitro* (Figure 2K). Re-expression of wild-type AAK1, but not kinase-dead AAK1, restored paclitaxel IC_50_ in AAK1 knockdown cells (Figure 2L), demonstrating that kinase activity is required for AAK1-mediated paclitaxel resistance. Notably, silencing clathrin or AP2M1—the canonical AAK1 substrate—did not sensitize cells to paclitaxel (Figure 2M), arguing against involvement of clathrin-dependent endocytosis. AAK1 silencing also did not affect multidrug resistance (MDR) activity or doxorubicin IC_50_ (Figures 2N–2O)^21,22^, collectively suggesting that AAK1 promotes paclitaxel resistance through an endocytosis- and efflux pump-independent mechanism.

### CDK1 phosphorylates AAK1 at T389 to enhance kinase activity upon paclitaxel treatment

Having established that AAK1 kinase activity is required for cellular adaptation to anti-tubulin agents, we next asked how paclitaxel treatment regulates AAK1. Interestingly, AAK1 activity was elevated following paclitaxel or nocodazole treatment (Figure 3A). Consistently, synchronization of cells with RO3306 followed by release revealed that AAK1 activity increased as cells entered mitosis (Figure 3B). These findings suggest that AAK1 is activated during mitotic arrest. To identify the upstream kinase responsible for AAK1 activation, we examined phosphorylation of AAK1 using phospho-substrate motif antibodies. We found that paclitaxel induced AAK1 phosphorylation with a CDK1 substrate motif, but not with motifs for PKA, AKT, ATM/ATR, or AMPK (Figure 3C). CDK1 inhibitor RO3306 blocked both paclitaxel-induced AAK1 phosphorylation and kinase activation (Figure 3D), and silencing CDK1, but not CDK2, also abrogated this effect (Figure 3E). *In vitro* kinase assays using a panel of CDKs and mitosis-related kinases confirmed that CDK1 directly phosphorylates AAK1 and increases its activity (Figure 3F). Proximity ligation assay (PLA) supported proximity between CDK1 and AAK1, which was enhanced by paclitaxel treatment (Figure 3G). Collectively these data suggest that CDK1 directly phosphorylates AAK1 to increase its kinase activity upon paclitaxel treatment.

**Figure 3.**
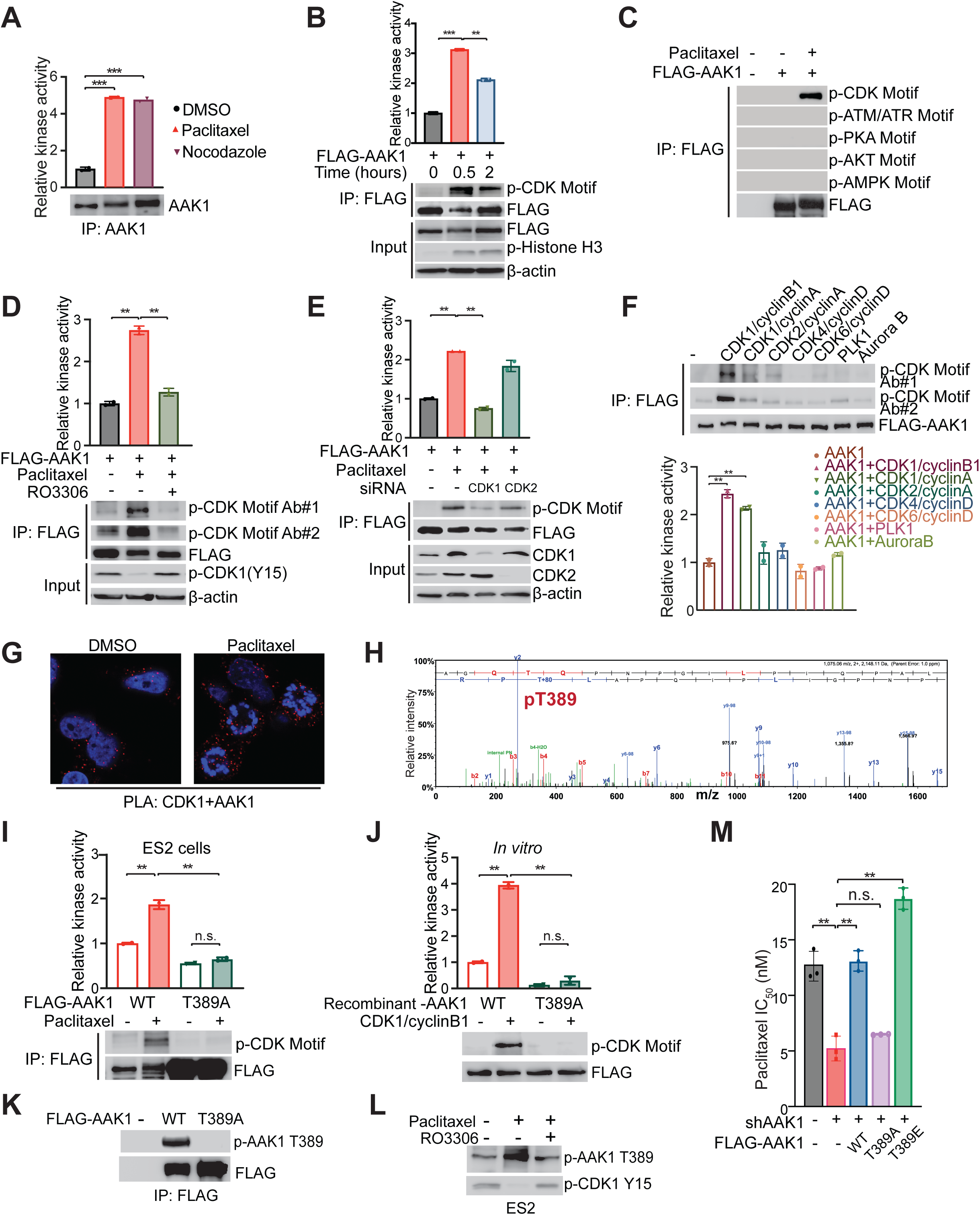
CDK1 phosphorylates AAK1 at T389 to promote its kinase activity and paclitaxel resistance. (A) AAK1 kinase activity in ES-2 cells after DMSO, paclitaxel or nocodazole treatment, with AAK1 immunoblot. (B) AAK1 kinase activity and p-CDK substrate motif signal after RO3306 synchronization and release (0, 0.5 and 2 h), with p-Histone H3 marking mitotic cells. (C) Phospho-substrate motif profiling of immunoprecipitated FLAG-AAK1, showing CDK substrate motif but not ATM/ATR, PKA, AKT or AMPK motif phosphorylation upon paclitaxel. (D) The CDK1 inhibitor RO3306 blocks paclitaxel-induced AAK1 phosphorylation and kinase activation; p-CDK1 (Y15) confirms CDK1 inhibition. (E) siRNA against CDK1, but not CDK2, abrogates paclitaxel-induced AAK1 phosphorylation and activation. (F) *In vitro* kinase assays with a panel of CDKs and mitotic kinases (CDK1/cyclin B1, CDK1/cyclin A, CDK2/cyclin A, CDK4/cyclin D, CDK6/cyclin D, PLK1 and Aurora B), showing that CDK1/cyclin B1 phosphorylates AAK1 on a CDK substrate motif and enhances its kinase activity (quantification below). (G) PLA showing that the CDK1-AAK1 interaction is enhanced by paclitaxel. (H) Phospho-mass spectrometry spectrum identifying AAK1 pT389. (I–J) T389A blunts paclitaxel-induced AAK1 activation in ES-2 cells (I) and *in vitro* with recombinant CDK1/cyclin B1 (J). (K) Validation of the AAK1 pT389 phospho-specific antibody on FLAG-AAK1 wild-type versus T389A. (L) RO3306 blocks paclitaxel-induced AAK1 T389 phosphorylation in ES-2 cells. (M) Paclitaxel IC_50_ in AAK1 knockdown cells reconstituted with FLAG-AAK1 wild-type, T389A or T389E. Data are mean ± SD. *p < 0.05, **p < 0.01, ***p < 0.001, n.s. not significant. (A-G, I-M) Results of one representative experiment from 2-3 independent experiments are shown.

We subsequently performed phospho-mass spectrometry analysis and identified T389 as the key CDK1-mediated phosphorylation site on AAK1 (Figure 3H). Furthermore, phospho-deficient AAK1 T389A mutant blunted paclitaxel-induced AAK1 phosphorylation and activation both in cells and *in vitro* (Figures 3I–3J). To further verify the functional significance of T389 phosphorylation, we developed an AAK1 pT389 phospho-specific antibody and confirmed that paclitaxel induces AAK1 T389 phosphorylation in a CDK1-dependent manner (Figures 3K–3L). Additionally, AAK1 wild-type and phospho-mimetic T389E, but not T389A, restored the paclitaxel IC_50_ in AAK1 knockdown cells (Figure 3M). Together, these data demonstrate that CDK1-mediated T389 phosphorylation enhances AAK1 kinase activity and promotes paclitaxel resistance.

### Caspase-7 cleavage at D767 generates an AAK1 fragment that supports paclitaxel tolerance

While examining AAK1 activation following paclitaxel treatment, we made a striking observation: in addition to the full-length AAK1 band (∼140 kDa), a faster-migrating band (∼100 kDa) appeared in paclitaxel-treated ovarian cancer cells (Figure 4A) and disappeared in AAK1 knockdown cells (Figure 4B). Although two AAK1 isoforms arising from alternative splicing have been reported in certain tissues^12^, RNA-seq analysis revealed no evidence of alternative splicing in ovarian cancer cells. Cycloheximide chase experiments showed that the short form progressively accumulated while full-length AAK1 correspondingly decreased upon paclitaxel treatment (Figure 4C), suggesting a post-translational mechanism. Formation of the truncated protein was abolished by the pan-caspase inhibitor Z-VAD but was unaffected by a proteasome inhibitor or by other classes of protease inhibitors (Figure 4D), implicating caspase-dependent proteolysis.

**Figure 4.**
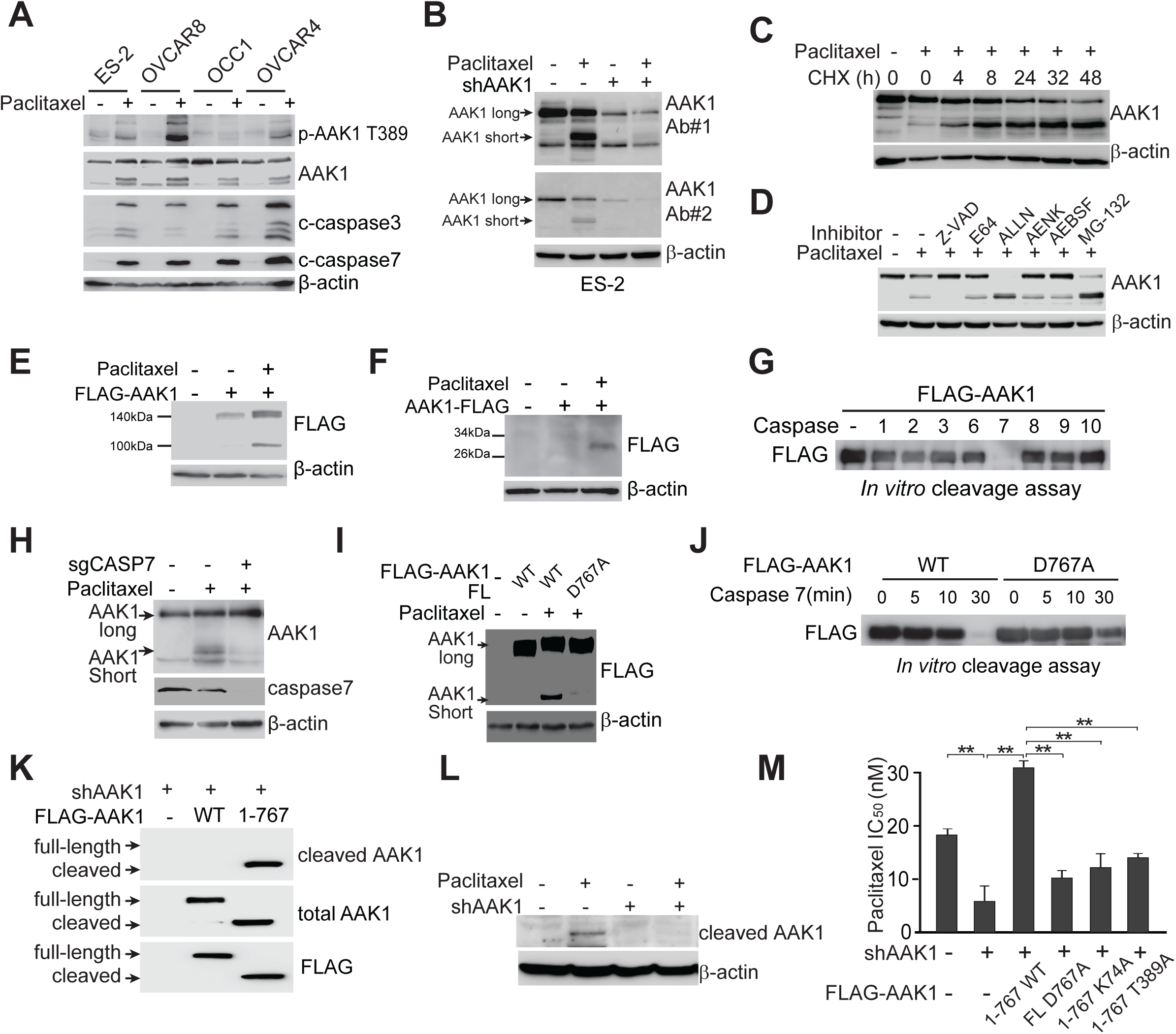
Caspase-7 cleaves AAK1 at D767 upon paclitaxel treatment. (A) Immunoblot of ES-2, OVCAR8, OCC1 and OVCAR4 cells with or without paclitaxel. (B) Immunoblot with two independent AAK1 antibodies in ES-2 cells, showing a paclitaxel-induced fast-migrating AAK1 species that is lost upon AAK1 knockdown. (C) Cycloheximide chase showing progressive accumulation of the short form during paclitaxel treatment. (D) The pan-caspase inhibitor Z-VAD, but not E64, ALLN, AENK, AEBSF or MG-132, blocks formation of the short form. (E-F) N-terminal FLAG-AAK1 yields a ∼100 kDa fragment (E) and C-terminal AAK1-FLAG a ∼26 kDa fragment (F) upon paclitaxel treatment. (G) *In vitro* cleavage of FLAG-AAK1 by a panel of recombinant caspases (caspases 1, 2, 3, 6, 7, 8, 9 and 10), identifying caspase-7 as the only protease that cleaves AAK1. (H) Paclitaxel-induced AAK1 cleavage is lost in sgCASP7 cells. (I) D767A blocks paclitaxel-induced cleavage in ES-2 cells. (J) D767A blocks caspase-7-mediated cleavage *in vitro* over a 30-min time course. (K) Validation of the AAK1 1-767 cleaved-form-specific monoclonal antibody in AAK1 knockdown cells reconstituted with FLAG-AAK1 wild-type or 1-767. (L) The cleaved-form-specific antibody detects endogenous paclitaxel-induced AAK1 1-767, which is lost upon AAK1 knockdown. (M) Paclitaxel IC_50_ in AAK1 knockdown cells reconstituted with AAK1 1-767, full-length D767A, 1-767 K74A or 1-767 T389A. Data are mean ± SD, n = 3. **p < 0.01. Results of one representative experiment from 2-3 independent experiments are shown.

To map the cleavage site, we expressed full-length AAK1 tagged with FLAG at either the N- or C-terminus. N-terminal FLAG-AAK1 generated a ∼100 kDa fragment, while C-terminal FLAG-AAK1 produced a ∼26 kDa fragment upon paclitaxel treatment (Figures 4E-4F), indicating cleavage near the C-terminus. Screening a panel of recombinant caspases (caspase 1, 2, 3, 6, 7, 8, 9 and 10) identified caspase-7 as the only tested protease that directly cleaves AAK1 under these *in vitro* conditions (Figure 4G). Consistently, paclitaxel treatment failed to induce AAK1 cleavage in caspase-7 knockout cells (Figure 4H), establishing caspase-7 as the key protease mediating AAK1 cleavage upon paclitaxel treatment.

To identify the precise cleavage site, we purified N-terminal FLAG-AAK1 from paclitaxel-treated cells and performed mass spectrometry analysis. A unique aspartic acid-ending peptide (SAGTAEKRKGGQTVD, corresponding to AAK1 S753-D767) was identified exclusively in the paclitaxel-treated sample. D-to-A mutagenesis confirmed that D767A largely blocked paclitaxel-induced cleavage in cells (Figure 4I) and caspase-7-mediated cleavage *in vitro* (Figure 4J). To facilitate detection of endogenous cleavage, we subsequently developed a recombinant rabbit monoclonal antibody specifically recognizing cleaved AAK1 (1-767) but not the full-length protein (Figure 4K), which confirmed paclitaxel-induced AAK1 cleavage at D767 (Figure 4L) in cells.

We next tested whether proteolytic processing contributes to the biological activity of AAK1 during paclitaxel treatment. Expression of the cleaved AAK1 (1-767) restored paclitaxel tolerance in AAK1-depleted cells, whereas the cleavage-deficient mutant (D767A) failed to rescue the phenotype (Figure 4M). Moreover, neither the kinase-dead (1-767 K74A) nor the phospho-deficient (1-767 T389A) form of the cleaved fragment restored paclitaxel tolerance (Figure 4M), indicating that cleaved AAK1 promotes paclitaxel resistance through a mechanism requiring both its kinase activity and T389 phosphorylation. Together, these findings demonstrate that paclitaxel induces caspase-7-mediated proteolytic conversion of AAK1, producing a functionally distinct kinase that mediates the adaptive response to mitotic stress.

### Proteolytic conversion rewires the AAK1 interactome from endocytosis toward the CCR4-NOT complex

Full-length AAK1 contains an N-terminal kinase domain and two consecutive clathrin-binding domains (CBDs) at the C-terminus (Figure 5A). Cleavage at D767 removes the second CBD, and accordingly, cleaved AAK1 1-767 showed reduced interaction with clathrin compared to full-length AAK1 (Figure 5B). To determine whether cleavage confers new molecular interactions, we compared the interactomes of full-length and cleaved AAK1 using complementary TurboID proximity labeling and FLAG pulldown-mass spectrometry approaches (Figure 5C). Gene ontology (GO-term) pathway analysis revealed that RNA binding was highly enriched in the cleaved AAK1 interactome but not in the full-length interactome (Figure 5D). Notably, several components of the CCR4-NOT complex—the major eukaryotic deadenylase that controls mRNA poly(A) tails and regulates mRNA decay and translation^3,23,24^—were identified as binding partners of cleaved AAK1 but not full-length AAK1 (Figure 5E).

**Figure 5.**
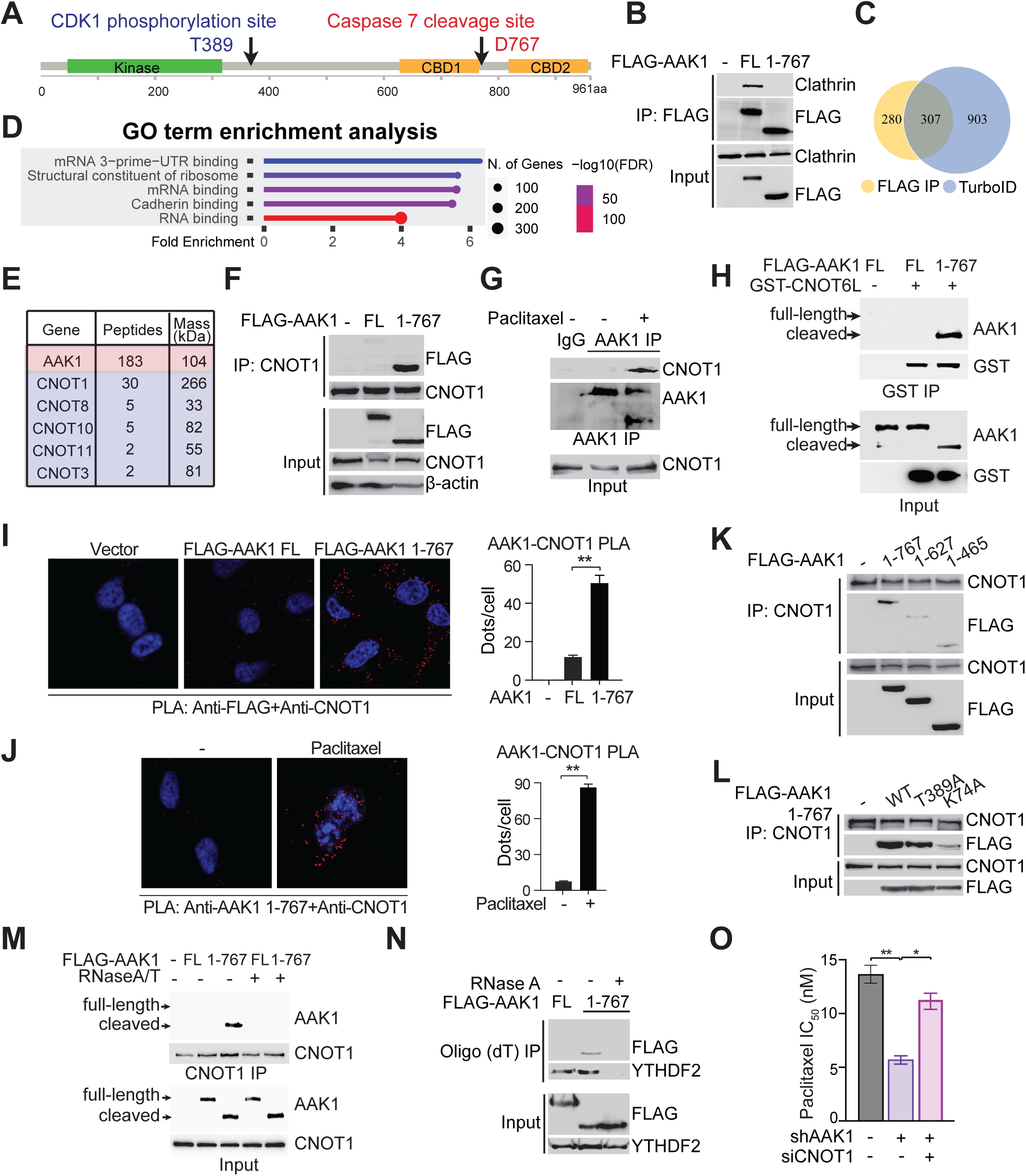
Proteolytic conversion redirects AAK1 from endocytosis to the CCR4-NOT deadenylase complex. (A) Domain organization of human AAK1 showing the kinase domain, the two clathrin-binding domains (CBD), the CDK1 site T389 and the caspase-7 cleavage site D767. (B) Co-immunoprecipitation showing reduced clathrin binding by AAK1 1-767 relative to full-length AAK1. (C) Overlap between proteins enriched with AAK1 1-767 by FLAG affinity purification-mass spectrometry and by TurboID proximity labeling. (D) GO-term enrichment of the AAK1 1-767 interactome, showing RNA binding and mRNA 3′UTR binding. (E) CCR4-NOT subunits identified in the AAK1 1-767 interactome, with peptide counts and molecular masses. (F) FLAG-AAK1 1-767 co-immunoprecipitates with endogenous CNOT1 far more efficiently than full-length AAK1. (G) Endogenous AAK1-CNOT1 co-immunoprecipitation is detected after paclitaxel treatment. (H) GST-CNOT6L pull-down showing binding of cleaved, but not full-length, AAK1. (I) PLA (anti-FLAG + anti-CNOT1) in cells expressing vector, FLAG-AAK1 FL or FLAG-AAK1 1-767, with quantification. (J) PLA using the AAK1 1-767-specific antibody together with anti-CNOT1, with or without paclitaxel, and quantification. (K) CNOT1 immunoprecipitation from cells expressing FLAG-AAK1 1-767, 1-627 or 1-465, showing that binding requires the region exposed by cleavage at D767. (L) CNOT1 immunoprecipitation from cells expressing FLAG-AAK1 1-767 wild-type, T389A or K74A, showing reduced binding by the kinase-dead mutant but not by T389A. (M) RNase A/T treatment reduces the AAK1 1-767-CNOT1 interaction. (N) Oligo(dT) pull-down showing AAK1 1-767, but not full-length AAK1, in complex with poly(A) mRNA; YTHDF2 serves as a positive control. (O) Paclitaxel IC_50_ in control and AAK1 knockdown cells with or without CNOT1 siRNA. Data are mean ± SD. *p < 0.05, **p < 0.01. (B, F-O) Results of one representative experiment from 2-3 independent experiments are shown.

We next validated these interactions biochemically and in cells. Cleaved AAK1 1-767 interacted substantially more strongly with CNOT1 than full-length AAK1 (Figure 5F), and endogenous association between AAK1 and CNOT1 was readily detected following paclitaxel treatment, when the cleaved AAK1 species is generated (Figure 5G). GST pull-down assays further demonstrated interaction between cleaved AAK1 and the catalytic CCR4-NOT subunit CNOT6L (Figure 5H). Proximity ligation assay (PLA) confirmed the preferential proximity between CNOT1 and cleaved AAK1 1-767 compared to full-length (Figure 5I). Importantly, using our AAK1 1-767 cleaved-form specific antibody, we verified by PLA that paclitaxel-induced endogenous AAK1 1-767 associates with CNOT1 *in situ* (Figure 5J). Further truncation of AAK1 to 1-627 or 1-465 substantially weakened the CNOT1 association (Figure 5K). The phospho-deficient T389A mutant showed slightly reduced binding to CNOT1 compared to wild-type, and the kinase-dead 1-767 K74A mutant showed strong reduction of CNOT1 binding (Figure 5L), indicating that catalytic activity contributes to complex formation. We next examined whether RNA participates in complex formation. RNase A/T treatment partially disrupted the interaction between cleaved AAK1 and CNOT1 (Figure 5M), suggesting that RNA contributes to stabilization of the AAK1-CCR4-NOT complex. Consistently, oligo(dT) pull-down assays showed that cleaved AAK1, but not full-length, forms a complex with mRNA (Figure 5N). Functionally, CNOT1 depletion partially reversed the decreased paclitaxel IC_50_ in AAK1 knockdown cells (Figure 5O), supporting a functional link between AAK1 1-767, the CCR4-NOT complex and paclitaxel sensitivity.

### Cleaved AAK1 suppresses CCR4-NOT-mediated mRNA decay to sustain metabolic adaptation during mitotic arrest

We next asked whether recruitment of AAK1 1-767 alters CCR4-NOT function during paclitaxel stress. Analysis of RNA targets revealed substantial overlap between AAK1 1-767-associated transcripts and CNOT3 targets, particularly under paclitaxel treatment (Figure 6A), suggesting that cleaved AAK1 and CCR4-NOT engage a shared RNA population. To determine whether the interaction of cleaved AAK1 with the CCR4-NOT complex influences its deadenylase activity, we used an MS2 tethering assay^25,26^. Tethered CNOT6L accelerated reporter mRNA degradation, and overexpression of cleaved AAK1 significantly inhibited this CNOT6L-mediated mRNA degradation (Figure 6B). Consistently, expression of AAK1 1-767, but not D767A, was associated with reduced deadenylase activity in cell-purified GST-CNOT6L preparations in AAK1 knockdown cells (Figure 6C). To examine whether AAK1 influences CCR4-NOT subunit association, we immunoprecipitated CNOT6 from paclitaxel-treated cells with or without AAK1 knockdown. AAK1 depletion increased the recovery of CNOT1 in CNOT6 immunoprecipitates (Figure 6D), suggesting that AAK1 alters CCR4-NOT subunit association under paclitaxel treatment.

**Figure 6.**
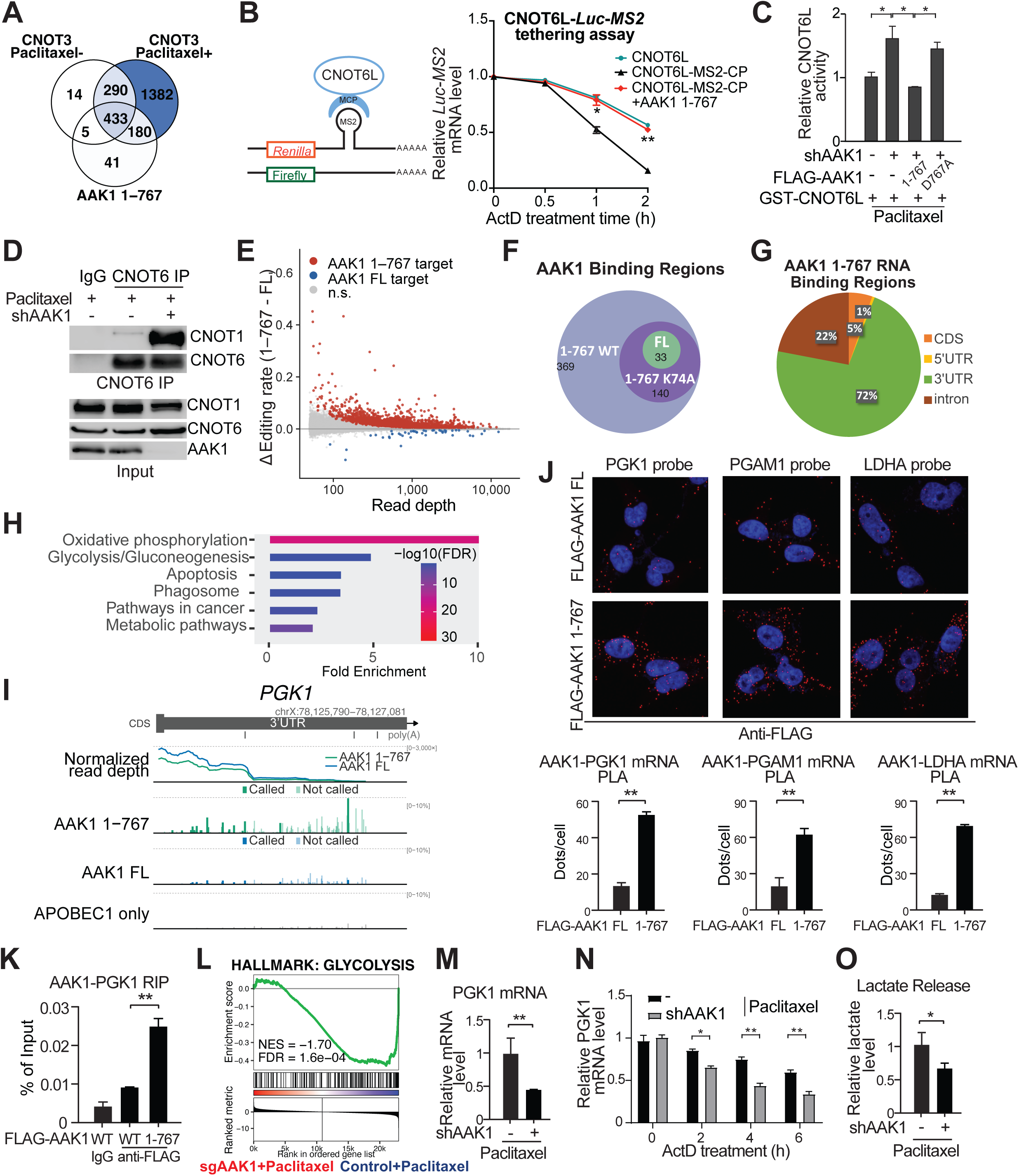
Cleaved AAK1 suppresses CCR4-NOT-mediated mRNA decay to promote glycolytic adaptation during mitotic arrest. (A) Overlap between AAK1 1–767-associated transcripts and CNOT3 targets in the presence or absence of paclitaxel, revealing a shared RNA population engaged by AAK1 1–767 and the CCR4-NOT complex. (B) MS2 tethering assay. Left, schematic of the Renilla-MS2 reporter and the Firefly control. Right, relative Luc-MS2 reporter mRNA over an actinomycin D time course in cells expressing CNOT6L, CNOT6L-MS2-CP, or CNOT6L-MS2-CP plus AAK1 1-767. (C) Relative deadenylase activity of GST-CNOT6L purified following paclitaxel treatment from control cells and AAK1-knockdown cells with no AAK1 rescue, FLAG-AAK1(1-767) rescue or FLAG-AAK1(D767A) rescue. GST-CNOT6L preparations were analyzed using the FRET assay. (D) CNOT6 was immunoprecipitated from paclitaxel-treated cells with or without AAK1 knockdown, and CNOT1 co-immunoprecipitation was assessed by immunoblotting. (E) Differential STAMP C-to-U editing between cells expressing an AAK1(1-767)-APOBEC1 fusion and cells expressing a full-length AAK1-APOBEC1 fusion. Each point is an edited cytosine (n = 186,144 sites passing a read-depth filter of ≥ 50 and a background editing rate < 0.05 in the APOBEC1-only control); the y axis is the difference in editing rate between the two fusions. Sites were called with a Cochran-Mantel-Haenszel test stratified by biological replicate (n = 3) using Benjamini-Hochberg-adjusted P < 0.05, common odds ratio ≥ 1.2 and |ΔE| ≥ 0.005 (red, sites edited more by AAK1 1-767; blue, sites edited more by full-length AAK1; grey, not significant). (F) Overlap of called RNA-binding regions among AAK1 1-767 wild-type (n = 545), kinase-dead AAK1 1-767 K74A (n = 174) and full-length AAK1 (FL, n = 38). Regions unique to K74A (n = 1), unique to FL (n = 2) and shared only by 1-767 and FL (n = 3) are too small to be drawn. (G) Distribution of AAK1 1-767 binding regions across transcript features. (H) Pathway enrichment of AAK1 1-767-bound transcripts. (I) Genome-browser view of the PGK1 locus showing normalized read depth and called editing sites for AAK1 1-767, AAK1 FL and the APOBEC1-only control. (J) RNA-protein PLA with PGK1, PGAM1 and LDHA probes in cells expressing FLAG-AAK1 FL or 1-767, with quantification below. (K) RIP-qPCR showing enhanced association of FLAG-AAK1 1-767 with PGK1 mRNA relative to full-length AAK1. (L) GSEA of RNA-seq data from control and AAK1 sgRNA cells treated with paclitaxel, showing suppression of the glycolysis hallmark signature. (M-O) PGK1 mRNA level, PGK1 mRNA decay after actinomycin D (N) and (O) lactate release in control and AAK1 knockdown cells treated with paclitaxel. Data are mean ± SD from three independent biological experiments unless otherwise indicated; two-tailed Student’s t-test. *p < 0.05, **p < 0.01. Results of one representative experiment from 2-3 independent experiments are shown.

Having established that AAK1 1-767 associates with CCR4-NOT and suppresses CNOT6L-mediated mRNA decay, we next sought to identify the endogenous transcripts engaged by cleaved AAK1 in an unbiased manner. We therefore performed Surveying Targets by APOBEC-Mediated Profiling (STAMP) coupled with whole-transcriptome sequencing^27^. Comparison of AAK1 1-767 with full-length AAK1 revealed a striking expansion of RNA engagement by the cleaved protein, with AAK1 1-767-specific editing events readily distinguishable from those associated with full-length AAK1 (Figure 6E). At the level of called RNA-associated regions, AAK1 1-767 is associated with 545 regions, compared with 38 for full-length AAK1. The kinase-dead AAK1 1-767 K74A mutant recovered 174 regions, indicating that kinase activity contributes to the expanded RNA interactome of AAK1 1-767 (Figure 6F). Mapping AAK1-associated regions revealed that the majority localized to 3′ untranslated regions (3′ UTRs) (Figure 6G), consistent with the established function of CCR4-NOT in deadenylating poly(A) tails and regulating mRNA stability. Pathway enrichment analysis of AAK1 1-767-interacting transcripts identified metabolic pathways including oxidative phosphorylation and glycolysis/gluconeogenesis (Figure 6H). Key glycolytic enzyme transcripts—PGK1, PGAM1, LDHA and LDHB—were among the most prominent targets (Figure 6I, Figures S1A-C), and inspection of individual loci revealed dense, AAK1 1-767-specific editing across the 3′ UTR, only sparse editing by full-length AAK1, and no called sites in the APOBEC1-only control (Figure 6I).

We next validated these candidate targets experimentally. RNA-protein proximity ligation assays demonstrated that cleaved AAK1, but not full-length AAK1, associated with the mRNAs of PGK1, PGAM1 and LDHA in situ (Figure 6J), supporting preferential association of cleaved AAK1 with these glycolytic transcripts. RNA immunoprecipitation followed by quantitative PCR confirmed significantly enhanced association of cleaved AAK1 with PGK1 mRNA compared with full-length AAK1 (Figure 6K). To determine whether these interactions influence gene expression during paclitaxel treatment, we performed RNA sequencing following AAK1 depletion. Gene set enrichment analysis identified glycolysis as one of the most significantly downregulated pathways in AAK1-deficient cells exposed to paclitaxel (Figure 6L). Consistent with this observation, depletion of AAK1 reduced PGK1 mRNA abundance, accelerated PGK1 mRNA decay and significantly decreased lactate release following paclitaxel treatment (Figures 6M-6O), indicating impaired glycolytic activity. Together, these findings support a model in which proteolytic conversion enables AAK1 to suppress CCR4-NOT-mediated mRNA decay and preserve glycolytic transcripts, contributing to metabolic adaptation during paclitaxel-induced mitotic arrest.

### Small molecule AAK1 inhibitors sensitize tumor models to paclitaxel

Resistance to anti-tubulin agents remains a major therapeutic challenge. Because AAK1 kinase activity is required for adaptation to paclitaxel-induced mitotic arrest, we next investigated whether pharmacological inhibition of AAK1 could sensitize tumor cells to paclitaxel. Given that AAK1 inhibitors have been developed for neuropathic pain with favorable safety profiles^13,14^ and one of these inhibitors has advanced to Phase 3 evaluation in diabetic peripheral neuropathic pain, we pursued a drug repurposing strategy. Four different small-molecule AAK1 inhibitors, at concentrations that did not affect cell viability alone, robustly sensitized ovarian cancer cells to paclitaxel (LP935509, LP922761, LX9211 and SGC-AAK1-1; Figure 7A). This paclitaxel-sensitizing effect was largely abolished in AAK1 knockdown cells (Figure 7B), supporting an AAK1-dependent component of the sensitizing effect. LP935509 combined with paclitaxel increased apoptosis beyond either agent alone (Figure 7C), and transcriptional profiling showed that the combination suppressed the glycolysis hallmark gene set (Figure 7D). LP935509 also sensitized a broad panel of ovarian cancer patient-derived organoids, primary cancer cells and cell lines, including paclitaxel-resistant sublines, to paclitaxel (Figure 7E). LP935509 significantly improved the paclitaxel response in an ovarian cancer cell line xenograft (Figure 7F), an ovarian cancer patient-derived xenograft (Figure 7H) and a head and neck cancer patient-derived xenograft (Figure 7I)^28,29^. No overt histopathological abnormalities were observed in the examined organs of combination-treated animals at endpoint (Figure 7G). Collectively, our data support repurposing clinically developed AAK1 inhibitors to enhance paclitaxel efficacy and provide a preclinical foundation for clinical evaluation of this combination strategy.

**Figure 7.**
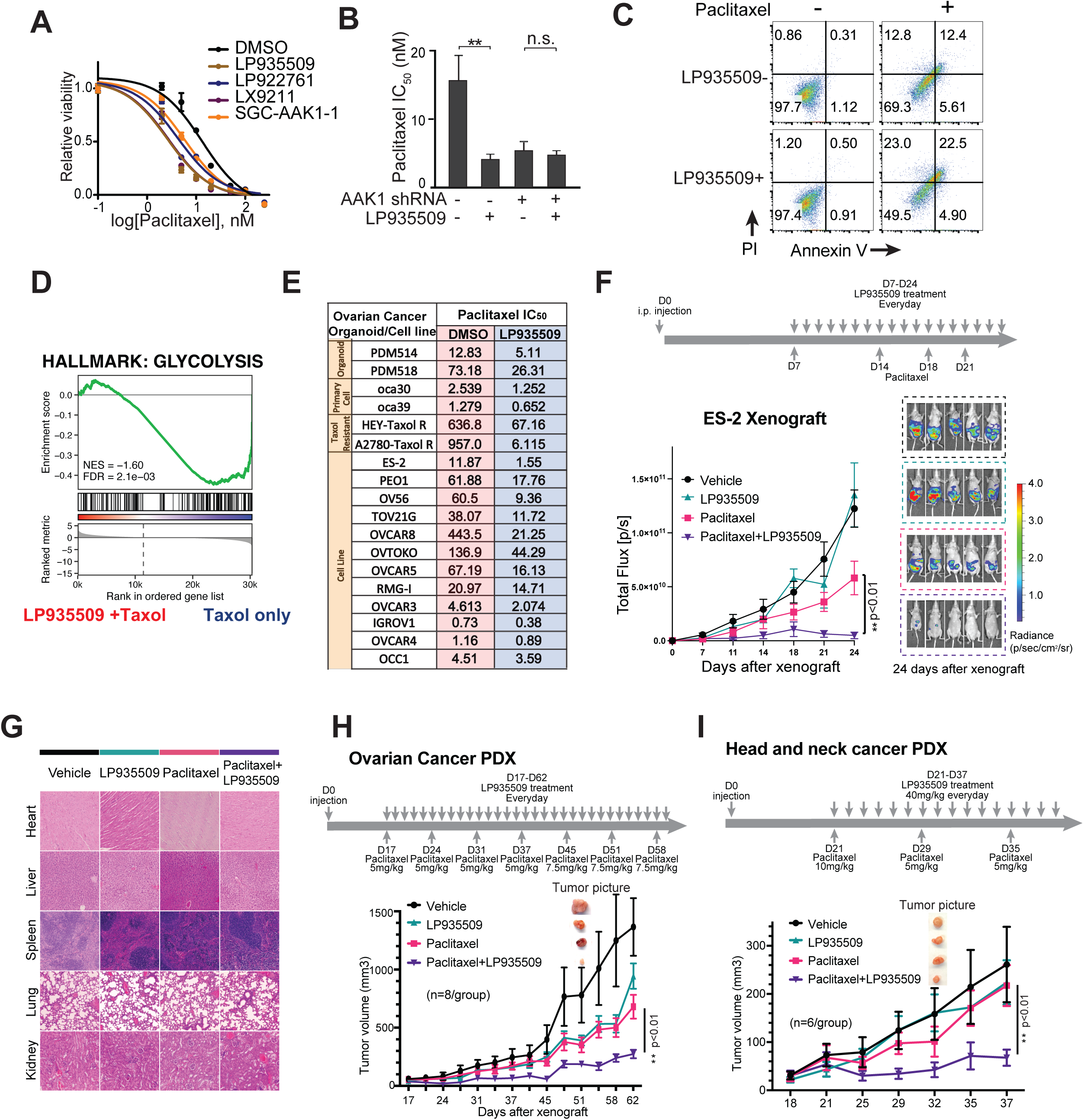
Small molecule AAK1 inhibitors sensitize ovarian and head and neck tumor models to paclitaxel *in vitro* and *in vivo*. (A) Paclitaxel dose-response curves for ES-2 cells co-treated with DMSO or one of four AAK1-targeting inhibitors (LP935509, LP922761, LX9211, SGC-AAK1-1) at non-cytotoxic concentrations. (B) Paclitaxel IC_50_ in control and AAK1 knockdown ES-2 cells with or without LP935509. (C) Annexin V/PI flow cytometry in ES-2 cells treated with paclitaxel, LP935509 or the combination. (D) GSEA of RNA-seq data from cells treated with LP935509 plus paclitaxel versus paclitaxel alone, showing suppression of the glycolysis hallmark signature. (E) Paclitaxel IC_50_ (nM) with DMSO or LP935509 across ovarian cancer patient-derived organoids, primary cancer cells, taxol-resistant sublines and established cell lines. (F) Treatment schedule, bioluminescence imaging and tumor growth curves of luciferase-expressing ES-2 xenografts treated with vehicle, LP935509, paclitaxel or the combination (n = 5 per group). (G) H&E organ histology of mice receiving vehicle, paclitaxel, LP935509 or combination as in (F). (H) Treatment schedule, tumor growth curves and representative tumors from ovarian cancer patient-derived xenografts (TM00335) treated with vehicle, LP935509, paclitaxel or the combination (n = 8 per group). (I) Treatment schedule, tumor growth curves and representative tumors from a head and neck cancer patient-derived xenograft (TM01145) treated with vehicle, LP935509, paclitaxel or the combination (n = 6 per group). Data are mean ± SD. **p < 0.01. (A-E) Results of one representative experiment from 2-3 independent experiments are shown.

## Discussion

During the prolonged mitotic arrest imposed by anti-tubulin chemotherapy, AAK1 is converted into a regulator of mRNA stability: CDK1-mediated phosphorylation at T389 enhances its kinase activity, caspase-7 cleavage at D767 removes the second clathrin-binding domain, and the resulting fragment shows reduced clathrin association and increased association with the CCR4-NOT deadenylase, suppresses mRNA decay, and maintains glycolytic gene expression associated with metabolic fitness during arrest (Figure 8). The two inputs are separable but both necessary. T389 phosphorylation is largely dispensable for CCR4-NOT binding yet required for taxane tolerance, whereas kinase activity promotes CCR4-NOT association and is required for taxane tolerance, suggesting separable assembly and output steps.

**Figure 8.**
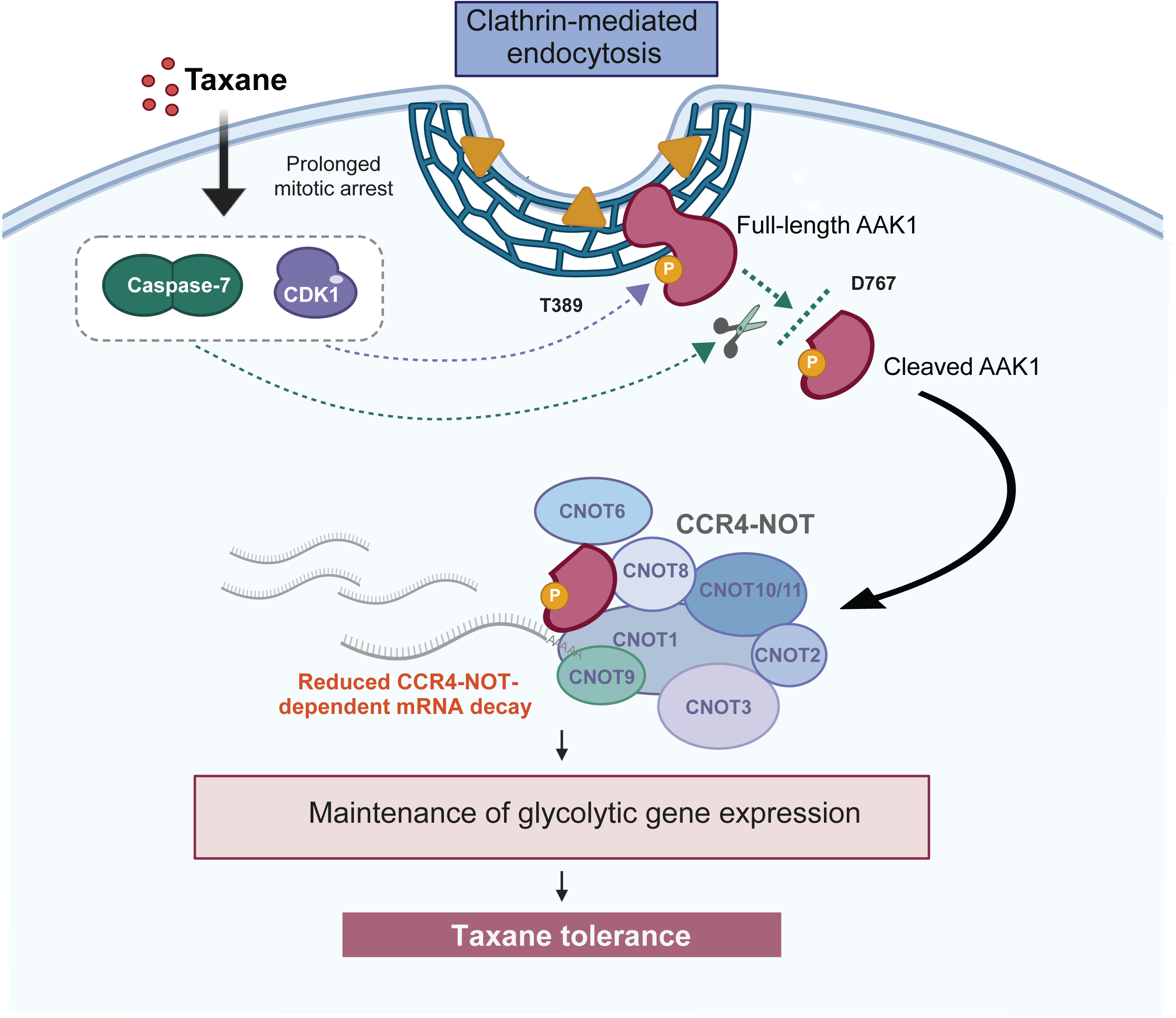
Working model of AAK1 conversion during mitotic arrest. During paclitaxel-induced mitotic arrest, CDK1 phosphorylation at T389 enhances AAK1 kinase activity, while caspase-7 cleavage at D767 removes the second clathrin-binding domain. The cleaved fragment shows reduced clathrin association and preferential association with CCR4–NOT. Genetic and biochemical findings support a model in which converted AAK1 suppresses CCR4–NOT-dependent mRNA decay, maintains glycolytic gene expression and contributes to paclitaxel tolerance.

Proteolytic cleavage is often viewed as an irreversible mechanism of protein inactivation during apoptosis, but cleavage can also generate fragments with distinct biological activities. AAK1 cleavage at D767 removes the second clathrin-binding domain, weakens association with the endocytic machinery, and generates a fragment that preferentially associates with a different protein complex. This functional conversion connects caspase-mediated proteolysis to post-transcriptional adaptation during chemotherapy-induced mitotic arrest. Consistent with this functional conversion, TurboID profiling showed strong enrichment of RNA-binding proteins in the AAK1 1-767 interactome, and STAMP revealed a markedly expanded set of RNA regions associated with AAK1 1-767 relative to full-length AAK1 (545 versus 38).

These findings also raise a broader question about the biological role of caspase-7. Caspase-7 is often grouped with caspase-3 as a largely redundant executioner because the two proteases show similar preferences toward short peptide substrates. However, studies using natural protein substrates demonstrate that caspase-3 and caspase-7 are biochemically and functionally distinct, with caspase-3 generally displaying broader substrate activity and caspase-7 exhibiting selectivity toward particular proteins^30^. This selectivity can arise from recognition mechanisms outside the catalytic pocket: a basic N-terminal exosite in caspase-7 promotes efficient cleavage of selected substrates, and the same region can bind RNA to facilitate proteolysis of RNA-binding proteins^31,32^. Caspase-7 can also support specialized stress responses rather than simply accelerating cellular demolition. For example, caspase-7-mediated cleavage and activation of acid sphingomyelinase promotes membrane repair after gasdermin or perforin pore formation, preserving cell integrity long enough for organized downstream responses to proceed^33^. Caspase-7 may therefore function as a selective proteome-remodeling protease.

Caspase-7 cleavage at D767 generates an active AAK1 fragment with a remodeled interactome and increased association with CCR4-NOT. Notably, the D767 cleavage site is embedded within QTVD↓, suggesting that determinants beyond a canonical DEVD-like tetrapeptide contribute to AAK1 recognition, and whether AAK1 is preferentially recognized by caspase-7 over caspase-3 *in vivo* remains to be established.

The functional connection between AAK1 1-767 and CCR4-NOT shows how proteolysis can convert a trafficking kinase into a regulator of mRNA fate, linking kinase signaling to post-transcriptional control. CCR4-NOT is a major eukaryotic deadenylase complex and a central regulator of mRNA turnover^3^. Our data support a model in which AAK1 1-767 suppresses CCR4-NOT-dependent mRNA decay and thereby preserves a subset of transcripts enriched for glycolytic enzymes. This connection is particularly relevant during paclitaxel-induced mitotic arrest, when glycolytic metabolism can support cellular survival^5,6^. By maintaining glycolytic gene expression, proteolytically converted AAK1 may provide the metabolic flexibility required for cells to withstand prolonged mitotic stress. AAK1 depletion also enhanced CNOT6-CNOT1 co-immunoprecipitation, suggesting that AAK1 can restrain association of the nuclease subunit with CNOT1-containing complexes. Together with the activity and RNA-decay assays, this finding supports a model in which altered CCR4-NOT subunit association contributes to AAK1-dependent control of mRNA turnover. The relevant kinase substrate remains unknown, and whether AAK1 acts through phosphorylation, RNA-dependent organization or direct competition for subunit association requires further study.

A very recent study implicated reduced deadenylation in global mRNA stabilization during prolonged mitotic arrest^4^. Our findings complement this work by linking AAK1 conversion to CCR4-NOT regulation, glycolytic gene expression, and taxane tolerance. Whether AAK1 contributes to the broader PABPC-dependent stabilization response remains to be established.

The therapeutic implications of these findings are strengthened by the availability of clinically developed AAK1 inhibitors. LX9211 (pilavapadin), for example, has completed Phase 2 and advanced to Phase 3 evaluation in diabetic peripheral neuropathic pain^14^, providing human pharmacology and tolerability experience for AAK1 inhibition. Across our *in vitro* and *ex vivo* models, multiple AAK1-targeting inhibitors enhanced paclitaxel activity in established cell lines, patient-derived organoids, primary tumor cells, and paclitaxel-resistant settings; *in vivo*, AAK1 inhibitor enhanced paclitaxel activity in an ovarian cancer cell line xenograft, an ovarian cancer patient-derived xenograft and a head and neck cancer patient-derived xenograft, indicating that the dependency is not confined to a single tumor type. Importantly, no overt histopathological abnormalities were observed in the examined organs of combination-treated animals at endpoint. Together, these findings provide a mechanistic and preclinical foundation for repurposing clinically developed AAK1 inhibitors to enhance paclitaxel efficacy. Because AAK1 inhibition has also been investigated as an analgesic approach, future studies should determine whether an oncology regimen could retain effects on neuropathic pain while enhancing antitumor activity; this potential dual benefit remains to be established experimentally.

Collectively, our findings identify AAK1 conversion by CDK1 phosphorylation and caspase-7 cleavage as a potential resistance mechanism linking mitotic arrest to CCR4-NOT-mediated mRNA regulation (Figure 8). They provide a mechanistic rationale for repurposing clinically developed AAK1 inhibitors to improve taxane responses and show how chemotherapy-induced arrest can change a kinase’s functional role.

## Methods

### Reagents and resources

Reagents and resources, including DESeq2^34^, MaxQuant^35^, STAR^36^ and SAILOR^27^, are listed in Supplementary Table 1.

### Cell lines

Human ovarian cancer cell lines (ES-2, OVCAR5, SKOV3, OVCAR8, OVCAR3, OVCAR4, PEO1, HEYA8, A2780, OV56, TOV21G, OVTOKO, RMG-I, IGROV1, OCC1), primary ovarian cancer cells oca30 and oca39, and KB-3-1 (and its paclitaxel-resistant derivative KB-3-1^TaxolR^) were obtained from ATCC or collaborators and maintained in DMEM or RPMI-1640 supplemented with 10% FBS and 1% penicillin/streptomycin at 37°C with 5% CO_2_. All cell lines were authenticated by short tandem repeat profiling and tested negative for mycoplasma.

### Patient-derived organoids and primary cells

Patient-derived organoids (PDOs) were obtained from ATCC. PDOs were maintained in Matrigel domes with defined organoid medium per vendor’s recommendation.

### Animal models

All animal experiments were performed in accordance with protocols approved by the Institutional Animal Care and Use Committee. Female nude mice and NSG mice (6–8 weeks old) were used for xenograft and PDX studies.

### Kinome-wide shRNA screening

The lentiviral shRNA library targeting 781 human kinase genes (4,518 clones) was used as previously described^9^. KB-3-1 paclitaxel-resistant human carcinoma cells (KB-3-1^TaxolR^) cells were transduced with lentivirus pool targeting each of the 781 human kinase and kinase-related genes, then treated with PBS, a sublethal dose of paclitaxel (20 ng/mL) or puromycin (0.5 μg/mL). Cell viability was determined after 3 days of treatment using CellTiter-Glo (Promega).

### Cell viability and IC50 determination

Cells were seeded in 96-well plates and treated with serial dilutions of paclitaxel, docetaxel, vincristine, cisplatin, or doxorubicin for 72 hours. Cell viability was measured using CellTiter-Glo (Promega). IC_50_ values were calculated using nonlinear regression analysis in GraphPad Prism.

### Western blotting and immunoprecipitation

Standard western blotting and co-immunoprecipitation were performed as previously described^9^. Primary antibodies included anti-AAK1 (commercial and in-house AAK1 pT389 and AAK1 1-767 specific antibodies), anti-CNOT1, anti-CDK1, anti-caspase-7, anti-cleaved caspase-3, anti-FLAG, and anti-β-actin.

### In vitro kinase assay

Recombinant CDK1/cyclin B and AAK1 proteins were incubated with ATP in kinase buffer at 30°C for 30 minutes. AAK1 phosphorylation was detected by western blotting using AAK1 pT389 antibody or phospho-substrate motif antibodies. AAK1 kinase activity was subsequently measured using a standard kinase assay with Micro 2 peptide substrate (KEEQSQITSQVTGQIGWR).

### In vitro caspase cleavage assay

Recombinant caspases were incubated with purified AAK1 protein in cleavage buffer at 37°C for 2 hours. Cleavage products were analyzed by western blotting using anti-AAK1 and anti-FLAG antibodies.

### FRET-based CNOT6L deadenylase activity assay

GST-CNOT6L was purified following paclitaxel treatment from control cells and from AAK1-knockdown cells with no AAK1 rescue or with AAK1(1-767) or AAK1(D767A) rescue. Deadenylase activity was measured using a FRET-based assay adapted from Katsumura et al. (2022)^37^. Purified GST-CNOT6L in 10 μL of deadenylase buffer containing 50 mM Tris-HCl (pH 7.5), 150 mM NaCl, 10% glycerol, 1 mM DTT, and 5 mM MgCl₂ was mixed with 10 μL of 5′-6-FAM-labeled poly(A)₁₂ RNA (2 μM) and incubated for 60 min at 37°C. Reactions were terminated by adding 5 μL of 20 μM 3′-TAMRA-labeled oligo(dT) DNA probe prepared in 1% SDS, 20 mM Tris-HCl (pH 8.0), and 0.5 mM EDTA. Following hybridization for 30 min at room temperature, fluorescence was measured at excitation/emission wavelengths of 485/520 nm to assess deadenylase activity.

### TurboID proximity labeling and mass spectrometry

AAK1 wild-type or 1-767 fused to TurboID was stably expressed in ovarian cancer cells. Biotin labeling was induced by addition of 50 μM biotin for 10 minutes. Biotinylated proteins were captured on streptavidin beads and analyzed by LC-MS/MS. Parallel FLAG pulldown-mass spectrometry was performed as a complementary approach.

### STAMP (Surveying Targets by APOBEC-Mediated Profiling)

STAMP was performed as previously described^27^. Briefly, C-to-U editing sites were identified to map protein-RNA interaction sites at nucleotide resolution. Full-length AAK1, AAK1(1-767) and AAK1 (1-767) K74A were fused to APOBEC1 and stably expressed in ES-2 cells; APOBEC1 alone served as the editing-background control. Three biological replicates were prepared per construct. Total RNA was extracted for whole-transcriptome sequencing. Reads were aligned to hg38 and per-site depth (DP) and alternate-allele counts (AD) were called on each strand. C-to-U positions with non-zero depth in every sample and mean depth above 50 were tested. For each position, edited and unedited counts were arranged as a 2 × 2 × 3 contingency array stratified by replicate and compared by Cochran–Mantel–Haenszel test (mantelhaen.test, R stats), with Benjamini–Hochberg adjustment. Editing rate was defined as the mean of AD/DP across replicates and ΔE as the difference between the two samples compared. Positions with adjusted P < 0.05, common odds ratio ≥ 1.2 (or ≤ 1/1.2) and |ΔE| ≥ 0.005 were called differentially edited. Positions whose editing rate exceeded 0.05 in the APOBEC1-only control were excluded, leaving 186,144 of 187,754 tested positions. Called sites were annotated to hg38 exons, introns, 5′ UTRs, coding sequence and 3′ UTRs, preserving strand.

### RNA-seq and GSEA analysis

Total RNA was extracted from cells with or without AAK1 depletion and paclitaxel treatment using RNeasy Mini Kit (Qiagen). Libraries were prepared and sequenced on Illumina NovaSeq. Differential expression analysis was performed using DESeq2, and Gene Set Enrichment Analysis (GSEA) was performed using the Broad Institute GSEA software.

### Tethering reporter assay

The MS2-CP tethering assay was performed as previously described^25,26^. Luciferase reporter mRNA carrying eight MS2-binding sites was co-expressed with MS2-CP-CNOT6L fusion protein, with or without AAK1 1-767 overexpression. Reporter mRNA abundance was measured at 0, 0.5, 1 and 2 h after actinomycin D addition.

### Metabolic assays

Lactate production was quantified using a lactate assay kit (Promega).

### mRNA decay assay

PGK1 mRNA decay was assessed in control and AAK1-knockdown cells treated with paclitaxel. Following actinomycin D addition, PGK1 mRNA abundance was measured at 0, 2, 4 and 6 h.

### RNA-protein proximity ligation assay

RNA-protein proximity ligation assays were used to assess association of FLAG-tagged full-length AAK1 or AAK1(1-767) with PGK1, PGAM1 and LDHA mRNAs. Sequence-specific probes are listed in Supplementary Table 2.

### RNA immunoprecipitation and qPCR

Association of FLAG-tagged full-length AAK1 or AAK1(1-767) with PGK1 mRNA was assessed by RNA immunoprecipitation followed by qPCR. Anti-FLAG immunoprecipitation was compared with an IgG control, and PGK1 RNA enrichment was expressed as a percentage of input.

### Xenograft and PDX studies

For cell line xenografts, luciferase-labeled ES-2 cells were implanted into female NSG mice by intraperitoneal injection. Seven days after implantation, mice were randomized into four treatment groups (vehicle, LP935509, paclitaxel, or combination). LP935509 was administered daily by intraperitoneal injection from day 7 through day 24. Paclitaxel was administered intraperitoneally on days 7, 14, 18, and 21. Tumor growth was monitored by weekly bioluminescence imaging and quantified as total photon flux.

For patient-derived xenograft (PDX) studies, freshly expanded TM00335 or TM01145 tumor fragments were implanted subcutaneously into female NSG mice. When tumors reached approximately 50–100 mm³, mice were randomized into four treatment groups (vehicle, LP935509, paclitaxel, or combination). LP935509 was administered by intraperitoneal injection at 40 mg/kg/day on days 17-62 for the ovarian PDX and days 21-37 for the head and neck PDX. In the ovarian PDX, paclitaxel was administered intraperitoneally at 5 mg/kg on days 17, 24, 31 and 37, and at 7.5 mg/kg on days 45, 51 and 58. In the head and neck PDX, paclitaxel was administered intraperitoneally at 10 mg/kg on day 21 and at 5 mg/kg on days 29 and 35. Control animals received vehicle according to the same schedule. Tumor volumes were measured twice weekly using calipers and calculated as (length × width²)/2. At the end of the study, tumors were excised and weighed. Organs were collected for histopathological analysis at endpoint.

### Histology

Formalin-fixed, paraffin-embedded tumor and organ sections were analyzed by H&E staining.

### Proximity ligation assay (PLA)

PLA was performed using the Duolink PLA kit (Sigma-Aldrich) according to the manufacturer’s instructions to visualize protein-protein interactions between CDK1 and AAK1, or AAK1 1-767 and CNOT1, in situ.

### MDR activity assay

MDR activity was measured using a fluorescent dye efflux assay as previously described^22^.

### Oligo(dT) pull-down

Cells were lysed, and poly(A)+ mRNA-protein complexes were captured using oligo(dT) magnetic beads. Associated proteins were eluted and analyzed by western blotting for AAK1 and CNOT1.

### Statistical analysis

Statistical analyses were performed using GraphPad Prism. Two-tailed unpaired Student’s t-test was used for two-group comparisons. One-way ANOVA with Tukey’s post hoc test was used for multiple group comparisons. STAMP uses replicate-stratified Cochran–Mantel–Haenszel testing. p < 0.05 was considered statistically significant. Sample sizes and replicates are indicated in figure legends.

### Correspondence and material requests

Further information and requests for resources and reagents should be directed to and will be fulfilled by the corresponding author, Lingtao Jin.

### Materials availability

All unique reagents generated in this study, including the AAK1 pT389 phospho-specific antibody and AAK1 1-767 cleaved-form specific antibody, are available from the corresponding author with a completed Materials Transfer Agreement.

## Data availability

RNA-seq and STAMP are publicly available as of the date of publication. Accession numbers are listed in Supplementary Table 1. Any additional information required to reanalyze the data reported in this paper is available from the corresponding author upon request.

## Code availability

This paper does not report original code.

## Acknowledgements

The MS2 tethering reporter construct was a gift from Dr. Gene Yeo of University of California, San Diego. This work was supported by the National Cancer Institute of the National Institutes of Health under award numbers R37CA249305 (L.J.), R01CA256482 (S.H.), R01CA269782 (L.J.), R01CA300241 (L.J.), R01CA299621 (L.J.); by NIH S10 grant S10OD036297 (E.W.); by the Mary Kay Ash Foundation Innovative/Translational Cancer Research Grant (L.J.); and by the IIMS-CTSA Pilot Project (L.J.). We acknowledge the Shared Resources of the Mays Cancer Center, supported by NCI Cancer Center Support Grant P30 CA054174. The content is solely the responsibility of the authors and does not necessarily represent the official views of the National Institutes of Health.

## Author contributions

Conceptualization, Y.J., S.K., and L.J.; Methodology, Y.J., K.L., P.N., L.S., S.K., and L.J.; Investigation, Y.J., K.L., P.N., L.S., S.K., and L.J.; Formal Analysis, Y.J., K.L., P.N., Z.Z., L.S., S.K., and L.J.; Resources, E.W., X.M., N.Z., G.H., L.Z.S., R.V., Z.Q., G.Z., S.H., L.S., S.K., and L.J.; Visualization, Y.J. and L.J.; Writing, Y.J. and L.J.; Supervision, L.J.; Project Administration, Y.J. and L.J.; Funding Acquisition, L.J.

