## Supplemental Information for "Caspase-7 cleavage repurposes AAK1 to suppress mRNA decay and promote tolerance to anti-tubulin agents"

##### Supplementary Table 1. Reagents and resources.

Reference numbers in this table correspond to the main manuscript.

| REAGENT or RESOURCE | SOURCE | IDENTIFIER |
|---|---|---|
| **Antibodies** | | |
| Rabbit polyclonal anti-AAK1 | Cell Signaling Technology | Cat# 79832; RRID: AB_2799942 |
| Rabbit monoclonal anti-AAK1 pT389 (phospho-specific) | Sino Biological | N/A |
| Rabbit monoclonal anti-AAK1 1-767 (cleaved-form specific) | Sino Biological | N/A |
| Rabbit anti-CNOT1 | Proteintech | Cat# 14276-1-AP; RRID: AB_10888627 |
| Mouse anti-CDK1 | Cell Signaling Technology | Cat# 9116; RRID: AB_2074795 |
| Rabbit anti-Caspase 7 | Cell Signaling Technology | Cat# 9492; RRID: AB_2228313 |
| Rabbit anti-Cleaved Caspase 3 | Cell Signaling Technology | Cat# 9661; RRID: AB_2341188 |
| Mouse anti-FLAG (M2) | Sigma-Aldrich | Cat# F1804; RRID: AB_262044 |
| Rabbit anti-Phospho-CDK Substrate Motif (1) | Cell Signaling Technology | Cat# 9477; RRID: AB_2714143 |
| Rabbit anti-Phospho-CDK Substrate Motif (2) | Cell Signaling Technology | Cat# 14371; RRID: AB_2798466 |
| Rabbit anti-Phospho-Histone H3 (Ser10) | Cell Signaling Technology | Cat# 9701; RRID: AB_331535 |
| Mouse anti-β-Actin | Sigma-Aldrich | Cat# A5316; RRID: AB_476743 |
| Rabbit anti-Clathrin Heavy Chain | Cell Signaling Technology | Cat# 4796; RRID: AB_10828486 |
| Mouse anti-AP2M1 | BD Biosciences | Cat# 611351; RRID: AB_398873 |
| Anti-PGK1 | Cell Signaling Technology | Cat# 63536; RRID: AB_3698329 |
| Anti-LDHA | Cell Signaling Technology | Cat# 3582; RRID: AB_2066887 |
| Anti-PGAM1 | Cell Signaling Technology | Cat# 12098; RRID: AB_2736922 |
| Anti-CNOT6 | Cell Signaling Technology | Cat# 13415; RRID: AB_2798214 |
| Anti-DCP1A (P-body marker) | Cell Signaling Technology | Cat# 15365; RRID: AB_2798738 |
| Anti-G3BP1 (stress granule marker) | Cell Signaling Technology | Cat# 61559; RRID: AB_2909406 |

| REAGENT or RESOURCE | SOURCE | IDENTIFIER |
|---|---|---|
| Rabbit anti-AAK1 (Ab#2) | Sigma-Aldrich | Cat# HPA017931; RRID: AB_1844406 |
| Rabbit monoclonal anti-AAK1 pT389 #2 (phospho-specific) | This paper | N/A |
| Rabbit anti-Phospho-ATM/ATR Substrate Motif | Cell Signaling Technology | Cat# 6966; RRID: AB_10949894 |
| Rabbit anti-Phospho-PKA Substrate Motif | Cell Signaling Technology | Cat# 9624; RRID: AB_331817 |
| Rabbit anti-Phospho-AKT Substrate Motif | Cell Signaling Technology | Cat# 9614; RRID: AB_331810 |
| Rabbit anti-Phospho-AMPK Substrate Motif | Cell Signaling Technology | Cat# 5759; RRID: AB_10949320 |
| Rabbit anti-Phospho-CDK1 (Tyr15) | Cell Signaling Technology | Cat# 9111; RRID: AB_331460 |
| Rabbit anti-Cleaved Caspase-7 | Cell Signaling Technology | Cat# 9491; RRID: AB_2068144 |
| Rabbit anti-YTHDF2 | Proteintech | Cat# 24744-1-AP; RRID: AB_2687435 |
| Anti-GST | Sigma-Aldrich | Cat# G7781; RRID: AB_259965 |
| **Bacterial and virus strains** | | |
| Lentiviral shRNA library targeting 781 human kinases (4,518 clones) | Horizon Discovery | Cat# RHS4884 |
| pLKO.1-shAAK1 #1 | Horizon Discovery | TRCN0000001942 |
| pLKO.1-shAAK1 #2 | Horizon Discovery | TRCN0000001946 |
| siRNA targeting CDK1 | Invitrogen | S464 |
| siRNA targeting CDK2 | Invitrogen | S204 |
| siRNA targeting CNOT1 | Invitrogen | S22842 |
| pLKO.1-shClathrin | Horizon Discovery | TRCN0000007982 |
| pLKO.1-shAP2M1 | Horizon Discovery | TRCN0000060241 |
| **Chemicals, peptides, and recombinant proteins** | | |
| Paclitaxel | Sigma-Aldrich | Cat# T7402 |
| Docetaxel | Sigma-Aldrich | Cat# 01885 |
| Vincristine | Sigma-Aldrich | Cat# V8879 |
| Cisplatin | Sigma-Aldrich | Cat# P4394 |
| Doxorubicin | Sigma-Aldrich | Cat# D1515 |
| Nocodazole | Sigma-Aldrich | Cat# M1404 |
| LP935509 (AAK1 inhibitor) | MedChemExpress | Cat# HY-117626 |
| LX9211 (AAK1 inhibitor) | MedChemExpress | Cat# HY-134829 |
| LP922761 (AAK1 inhibitor) | MedChemExpress | Cat# HY-120179 |
| SGC-AAK1-1 (AAK1 inhibitor) | MedChemExpress | Cat# HY-123940 |
| RO3306 (CDK1 inhibitor) | Sigma-Aldrich | Cat# SML0569 |
| Z-VAD-FMK (pan-caspase inhibitor) | R&D Systems | Cat# FMK001 |
| MG132 (proteasome inhibitor) | Sigma-Aldrich | Cat# M7449 |
| Cycloheximide | Sigma-Aldrich | Cat# C7698 |

| REAGENT or RESOURCE | SOURCE | IDENTIFIER |
|---|---|---|
| Actinomycin D | Sigma-Aldrich | Cat# A1410 |
| Biotin (for TurboID) | Sigma-Aldrich | Cat# B4639 |
| Recombinant human Caspase 7 | R&D Systems | Cat# 823-C7 |
| Recombinant CDK1/Cyclin B | Invitrogen | Cat# PV3292 |
| Recombinant AAK1 (full-length) | This paper | N/A |
| Recombinant AAK1 T389A | This paper | N/A |
| Recombinant AAK1 T389E | This paper | N/A |
| Recombinant AAK1 1-767 | This paper | N/A |
| Recombinant human caspases 1, 2, 3, 6, 8, 9 and 10 | Enzo | Cat# ALX-850-243 |
| Recombinant CDK1/cyclin A, CDK2/cyclin A, CDK4/cyclin D, CDK6/cyclin D, PLK1, Aurora B | SignalChem | Cat# C22-18G (CDK1/cyclin A2); C29-10G (CDK2/cyclin A2); C31-10G (CDK4/cyclin D1); C35-18H (CDK6/cyclin D1); P41-10H (PLK1); A31-10G (Aurora B) |
| Recombinant GST-CNOT6L | This paper | N/A |
| E64 (cysteine protease inhibitor) | Sigma-Aldrich | Cat# E3132 |
| ALLN (calpain inhibitor) | Sigma-Aldrich | Cat# A6185 |
| AEBSF (serine protease inhibitor) | Sigma-Aldrich | Cat# A8456 |
| AENK (asparagine endopeptidase inhibitor) | Sigma-Aldrich | Cat# 533796 |
| **Critical commercial assays** | | |
| CellTiter-Glo Luminescent Cell Viability Assay | Promega | Cat# G7570 |
| Annexin V-FITC/PI Apoptosis Kit | Biolegend | Cat# 640914 |
| Duolink In Situ PLA Kit | Sigma-Aldrich | Cat# DUO92101 |
| Pierce Magnetic RNA-Protein Pull-Down Kit | Thermo Scientific | Cat# 20164 |
| MDR Activity Assay | Thermo Scientific | Cat# V13180 |
| Lactate Assay Kit | Promega | Cat# J5021 |
| RNeasy Mini Kit | Qiagen | Cat# 74104 |
| **Deposited data** | | |
| RNA-seq data (AAK1 knockdown ± paclitaxel) | This paper | GSE354511 |
| STAMP sequencing data | This paper | GSE354512; GSE354513 |
| **Experimental models: Cell lines** | | |
| Human: ES-2 (ovarian clear cell carcinoma) | ATCC | Cat# CRL-1978; RRID: CVCL_3509 |
| Human: OVCAR8 (ovarian high-grade serous) | NCI | RRID: CVCL_1629 |
| Human: PEO1 (ovarian high-grade serous) | Sigma-Aldrich | Cat# 10032308; RRID: CVCL_2686 |
| Human: HEYA8 (ovarian carcinoma) | ATCC | RRID: CVCL_8878 |
| Human: A2780 (ovarian carcinoma) | Sigma-Aldrich | Cat# 93112519; RRID: CVCL_0134 |
| Human: KB-3-1 (cervical carcinoma) | This paper | RRID: CVCL_2088 |
| Human: KB-3-1$^{TaxolR}$ (paclitaxel-resistant) | This paper | N/A |

| REAGENT or RESOURCE | SOURCE | IDENTIFIER |
|---|---|---|
| Human: HOSEpiC (primary ovarian surface epithelial) | ScienCell | Cat# 7310 |
| Human: OVCAR5 (ovarian carcinoma) | Sigma-Aldrich | Cat# SCC259; RRID: CVCL_1628 |
| Human: SKOV3 (ovarian adenocarcinoma) | ATCC | Cat# HTB-77; RRID: CVCL_0532 |
| Human: OVCAR3 (ovarian adenocarcinoma) | ATCC | Cat# HTB-161; RRID: CVCL_0465 |
| Human: OVCAR4 (ovarian carcinoma) | ATCC | RRID: CVCL_1627 |
| Human: OV56 (ovarian adenocarcinoma) | Sigma-Aldrich | Cat# 96020759; RRID: CVCL_2673 |
| Human: TOV21G (ovarian clear cell carcinoma) | ATCC | Cat# CRL-11730; RRID: CVCL_3613 |
| Human: OVTOKO (ovarian clear cell carcinoma) | JCRB Cell Bank | Cat# JCRB1048; RRID: CVCL_3117 |
| Human: RMG-I (ovarian clear cell carcinoma) | JCRB Cell Bank | Cat# JCRB0172; RRID: CVCL_1662 |
| Human: IGROV1 (ovarian carcinoma) | Sigma-Aldrich | Cat# SCC203; RRID: CVCL_1304 |
| Human: OCC1 (ovarian carcinoma) | ATCC | RRID: CVCL_C829 |
| Human: HEY-Taxol R (paclitaxel-resistant) | This paper | N/A |
| Human: A2780-Taxol R (paclitaxel-resistant) | This paper | N/A |
| Patient-derived organoid PDM514 | ATCC | Cat# PDM514; |
| Patient-derived organoid PDM518 | ATCC | Cat# PDM518; |
| Primary ovarian cancer cells oca30 | This paper | N/A |
| Primary ovarian cancer cells oca39 | This paper | N/A |
| **Experimental models: Organisms/strains** | | |
| Mouse: NSG (NOD.Cg-Prkdcscid Il2rgtm1Wjl/SzJ) | The Jackson Laboratory | JAX: 005557; RRID: IMSR_JAX:005557 |
| Mouse: NU/J | The Jackson Laboratory | JAX: 002019; RRID: IMSR_JAX:002019 |
| Ovarian cancer patient-derived xenograft | The Jackson Laboratory | Cat# TM00335 |
| Head and neck cancer patient-derived xenograft | The Jackson Laboratory | Cat# TM01145 |
| **Oligonucleotides** | | |
| qPCR primers for PGK1 | This paper | See Supplementary Table 2 |
| PLA oligo probes for PGK1, LDHA, PGAM1 | This paper | See Supplementary Table 2 |
| sgRNA targeting human AAK1 | This paper | See Supplementary Table 2 |
| **Recombinant DNA** | | |
| pLenti-FLAG-AAK1 wild-type (shRNA-resistant) | This paper | N/A |
| pLenti-FLAG-AAK1 K74A(kinase-dead) | This paper | N/A |
| pLenti-FLAG-AAK1 T389A (phospho-deficient) | This paper | N/A |
| pLenti-FLAG-AAK1 T389E (phospho-mimetic) | This paper | N/A |
| pLenti-FLAG-AAK1 D767A (cleavage-deficient) | This paper | N/A |
| pLenti-FLAG-AAK1 T765A | This paper | N/A |
| pLenti-FLAG-AAK1 T765E | This paper | N/A |
| pLenti-FLAG-AAK1 1-627, 1-465 and 1-767 | This paper | N/A |

| REAGENT or RESOURCE | SOURCE | IDENTIFIER |
|---|---|---|
| pLenti-FLAG-AAK1 1-767 K74A (kinase-dead) | This paper | N/A |
| pLenti-FLAG-AAK1 1-767 T389A | This paper | N/A |
| pLenti-N-FLAG-AAK1 (N-terminal tag) | This paper | N/A |
| pLenti-AAK1-FLAG-C (C-terminal tag) | This paper | N/A |
| pLenti-TurboID-AAK1 wild-type | This paper | N/A |
| pLenti-TurboID-AAK1 1-767 | This paper | N/A |
| pLenti-APOBEC1-AAK1 wild-type (for STAMP) | This paper | N/A |
| pLenti-APOBEC1-AAK1 1-767 and 1-767 K74A (for STAMP) | This paper | N/A |
| pLenti-MS2-CP-CNOT6L (tethering assay) | This paper | N/A |
| GST-AAK1 fragments (for pull-down) | This paper | N/A |
| GST-CNOT1 fragments (for pull-down) | This paper | N/A |
| FLAG-CNOT1 fragments (for co-IP mapping) | This paper | N/A |
| **Software and algorithms** | | |
| GraphPad Prism (v10) | GraphPad | https://www.graphpad.com; RRID: SCR_002798 |
| DESeq2 | Love et al.[34] | https://bioconductor.org/packages/DESeq2/; RRID: SCR_015687 |
| GSEA (Gene Set Enrichment Analysis) | Broad Institute | https://www.gsea-msigdb.org; RRID: SCR_003199 |
| ImageJ | NIH | https://imagej.net/ij/; RRID: SCR_003070 (ImageJ) |
| MaxQuant (mass spectrometry) | Cox and Mann[35] | https://www.maxquant.org; RRID: SCR_014485 |
| STAR (RNA-seq alignment) | Dobin et al.[36] | https://github.com/alexdobin/STAR; RRID: SCR_004463 |
| SAILOR (STAMP analysis) | Brannan et al.[27] | https://github.com/YeoLab/sailor |
| R (stats package; mantelhaen.test) | R Core Team | https://www.r-project.org; RRID: SCR_001905 |
| **Other** | | |
| Streptavidin magnetic beads | Thermo Fisher | Cat# 88816 |
| Oligo(dT) magnetic beads | Thermo Fisher | Cat# 61002 |
| Matrigel Basement Membrane Matrix | Corning | Cat# 354234 |
| Propidium Iodide | Sigma-Aldrich | Cat# P4170 |

#### Supplementary Table 2. Oligonucleotides used in this study.

| Oligonucleotide | Sequence (5′→3′) |
|---|---|
| ***RNA–protein proximity ligation assay probes*** | |
| PGK1_probe1 | GAGUAGUGAAACAGUGACAAAGCUAAUCAU |
| PGK1_probe2 | UUUUAUUGUUUACUUAAUGGAUCAUCAAUU |
| PGAM1_probe1 | GCAUUUCACUGCCAUGAAACAAGGCAGGAA |
| LDHA_probe1 | CACACGGUAAACAUCCACCUGGCUCAAGGG |
| PFKP_probe1 | AUACGUGCAUAAAGUGCUGAUAAAAUAACU |
| ***CRISPR–Cas9 single-guide RNA cloning oligonucleotides*** | |
| AAK1CRISPR-1-F | CACCGTGGCAAAATCATCACTACGA |
| AAK1CRISPR-1-R | AAACTCGTAGTGATGATTTTGCCAC |

Proximity ligation assay probes were used to detect the indicated endogenous transcripts in proximity to FLAG-tagged full-length or cleaved AAK1 (1-767) by RNA–protein proximity ligation assay (Figure 6).

CRISPR–Cas9 oligonucleotides were annealed and ligated into the BsmBI-digested lentiCRISPRv2 vector. The forward and reverse oligonucleotides carry CACCG and AAAC adapters, respectively, and encode the AAK1 protospacer 5′-TGGCAAAATCATCACTACGA-3′ preceded by a guanosine for U6-driven transcription.

### Figure S1

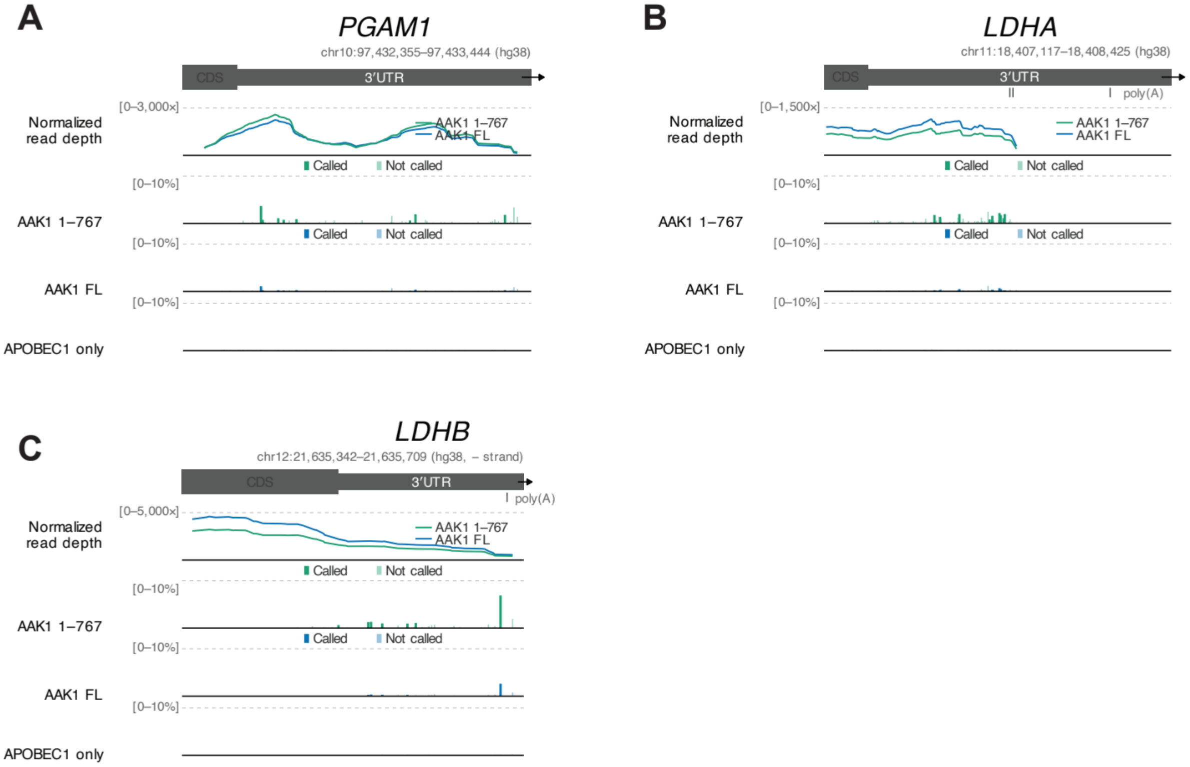


**Supplementary Figure 1. STAMP editing at additional glycolytic loci, related to Figure 6.** (A-C) Genome-browser views of the PGAM1 (A), LDHA (B) and LDHB (C) loci showing normalized read depth and called editing sites for AAK1 1-767, AAK1 FL and the APOBEC1-only control.
